# Visualizing drug inhibitor binding interactions using microcrystal electron diffraction

**DOI:** 10.1101/2020.04.27.064188

**Authors:** Max T.B. Clabbers, S. Zoë Fisher, Mathieu Coinçon, Xiaodong Zou, Hongyi Xu

## Abstract

Visualizing drug inhibitor binding interactions at the atomic level is important for both structure-based drug design and fragment-based screening methods. Rapid and uniform soaking with potentially less lattice defects make small macromolecular crystals attractive targets for studying ligand biding using 3D microcrystal electron diffraction (MicroED). However, so far no drug inhibitor binding interactions could unambiguously be resolved by electron diffraction. Here, we use MicroED to study the binding of a sulfonamide inhibitor to human carbonic anhydrase isoform II (HCA II). We show that MicroED data can efficiently be collected in-house on a conventional TEM from thin hydrated microcrystals after a brief soaking with the clinical drug inhibitor acetazolamide (AZM). The data are of high enough quality to unequivocally fit and resolve the inhibitor bound to the active site of the protein. We anticipate MicroED can play an important role in future drug discovery experiments, complementing existing methods in structural biology such as x-ray and neutron diffraction.

## Introduction

Small three-dimensional crystals are highly suitable for structure determination by electron diffraction, commonly referred to as microcrystal electron diffraction (MicroED) (Shi *et al.*, 2013; Nannenga & Gonen, 2019) and 3D electron diffraction (3D ED) (Gemmi *et al.*, 2019). MicroED data can efficiently be collected in-house on a conventional transmission electron microscope (TEM) using the rotation method (Nederlof *et al.*, 2013; Nannenga, Shi, Leslie *et al.*, 2014) from thin hydrated macromolecular crystals. Furthermore, small macromolecular crystals may have less defects and lower mosaicity than larger crystals, and any external changes such as ligand soaking or rapid flash-cooling can be applied faster and more uniformly (Cusack *et al.*, 1998; Wolff *et al.*, 2020). Small crystal volumes also have their disadvantages, notably the overall diffracting intensity is much weaker, and radiation damage is limiting the maximum attainable resolution and affecting data and model quality (Hattne *et al.*, 2018). In recent years, MicroED has emerged as a promising method for structural biology, determining structures of several known macromolecules (Shi *et al.*, 2013; Nannenga, Shi, Leslie *et al.*, 2014; Nannenga, Shi, Hattne *et al.*, 2014; Yonekura *et al.*, 2015; Clabbers *et al.*, 2017; de la Cruz *et al.*, 2017; Xu *et al.*, 2018; Nannenga & Gonen, 2019), and even solving a previously unknown protein structure using MicroED data (Xu *et al.*, 2019). These results illustrate how MicroED can complement existing methods in structural biology, loosening the size limitations imposed on the sample as (sub-)micron sized 3D crystals resist structure determination by x-ray and especially neutron diffraction (Henderson, 1995). Furthermore, biomolecules of low molecular weight that are still challenging for single particle cryo-EM (Henderson, 1995; Fan *et al.*, 2019) can be studied by MicroED.

An important application of structural biology is structure-based drug discovery and design, as it relies on detailed structural knowledge of the protein active site and the underlying molecular interactions of inhibitor binding (Blundell *et al.*, 2002). Similarly, high-throughput screening of a large number of fragments for possible interactions with the target protein can reveal binding interactions that inform the design of novel inhibitors (Collins *et al.*, 2018). The potential of MicroED for drug discovery was first indicated in recent work studying the binding of an inhibitor bevirimat (BVM) to HIV-1 viral protease (Purdy *et al.*, 2018). The structural model provided insight into the underlying interactions in inhibitor binding, but a unique binding pose could not be resolved from the MicroED data. We previously presented the structure of a novel R2lox ligand-binding metalloenzyme (Xu *et al.*, 2019). Although the ligand could not be resolved from the MicroED data, the substrate-binding pocket was reshaped and had an altered projected electrostatic contact potential distribution, indicating a different ligand binding interaction compared to structural homologues.

Building upon earlier results, we use MicroED to investigate drug inhibitor binding interactions to the active site of human carbonic anhydrase isoform II (HCA II), a small ubiquitous metalloenzyme with a molecular weight of 29 kDa. Carbonic anhydrases (CAs) catalyze the reversible reaction of CO_2_ hydration to produce HCO_3_^−^ and H^+^. The reaction has two components: CO_2_ hydration, and the rate-limiting proton transfer step. After the first half-reaction of CO_2_ hydration, a zinc-bound water molecule (ZW) is left bound to the active site metal (Zn). In the next step, the ZW is deprotonated to OH^−^, ready for a nucleophilic attack on the next incoming CO_2_ molecule. The generated proton (H^+^) of the ZW deprotonation is transported to the bulk solvent via a hydrogen-bonded water network and an internal proton shuttling residue, His64 (Coleman, 1967; Silverman & Mckenna, 2007). There are 15 expressed human CA isoforms (HCAs) showing some diversity in physiological and subcellular distribution. Of these, HCA II is found in red blood cells and has the highest activity of any HCAs with a k_cat_ of 10^−6^.s^−1^ and a k_cat_/K_M_ approaching the diffusion limit (10^8^ M^−1^.s^−1^) (Khalifah, 1971). The structural model of the native water-bound HCA II has been well characterized in the past, both by x-ray and neutron diffraction (Liljas *et al.*, 1972; Zöe Fisher *et al.*, 2010).

There are many known small molecular inhibitors of CA, from small anions to the widely prescribed sulfonamide-based inhibitors used to treat glaucoma, altitude sickness, and congestive heart failure (Coleman, 1967; Supuran *et al.*, 2004). Several crystal structures of CA complexes with sulfonamide inhibitors show binding interactions that can effectively shut down the enzyme: the ionized -NH group binds directly to the Zn; the -NH group donates a hydrogen bond to the hydroxyl group of Thr199 side chain; the sulfonamide O interacts with the amide backbone of Thr199; and finally binding of the inhibitor displaces the ZW (Sippel *et al.*, 2009; Fisher *et al.*, 2012). Neutron studies of HCA II in complex with various sulfonamides also revealed additional hydrogen bonding interactions and water displacements in the active site that are important determinants in understanding inhibitor binding (Fisher *et al.*, 2012; Aggarwal *et al.*, 2016; Kovalevsky *et al.*, 2018).

Here we present the structure of HCA II in complex with the inhibitor acetazolamide (AZM) demonstrating MicroED data are of sufficient quality to fit and resolve drug inhibitor binding. We demonstrate that drug binding can be studied efficiently by briefly soaking the 3D microcrystals with inhibitor and collecting MicroED data using a conventional TEM, making it effectively feasible to screen quickly for possible protein-inhibitor binding interactions on a home source. Data processing, structure determination, and fitting and refining the inhibitor bound to the active site was feasible using standard crystallographic routines. We validate the structural model by comparison with the native structure as a negative control to confirm the correct interpretation for the model of HCA II with the bound drug inhibitor determined from the MicroED data. Furthermore, we compare our MicroED model and data with previously determined x-ray and neutron crystal structures of the HCA II:AZM complex.

## Results

### Data collection

Inhibitor-bound protein complexes were obtained after 20 minutes of soaking HCA II microcrystals with the AZM inhibitor. MicroED data were collected using continuous rotation on both the native and inhibitor-bound HCA II crystals to 2.5 and 2.3 Å resolution, respectively (Fig. 1). In order to minimize the data quality being affected by radiation damage (Hattne *et al.*, 2018; Bücker *et al.*, 2020), data from small angular ranges of on average 10-30° with an accumulated electron dose of typically 2.0-6.0 e^−^/Å^2^ per dataset were used for structure analysis. For both native and ligand bound protein, merged data from 16 crystal datasets were used for structure determination, covering a combined angular range of −60 to +60° (Table 1). The weighted average of the unit cell parameters are close to values described in literature (Sippel *et al.*, 2009; Fisher *et al.*, 2012). Owing to preferred crystal orientation, and limited tilt range of the TEM goniometer stage, data completeness did not increase beyond about 73%.

**Table 1.** Data collection and refinement statistics

|  | HCA II * | HCA II:AZM * |
| --- | --- | --- |
| **Data collection** |  |  |
| Space group | $P2_1$ | $P2_1$ |
| Cell dimensions |  |  |
| $a, b, c$ (Å) | 42.52(1), 41.39(1), 72.80(2) | 42.53(2), 41.81(1), 72.24(3) |
| $\alpha, \beta, \gamma$ (°) | 90.00, 104.56(3), 90.00 | 90.00, 104.46(3), 90.00 |
| Resolution (Å) | 29.18-2.50 (2.60-2.50) | 35.89-2.30 (2.38-2.30) |
| $R_{\text{merge}}$ | 0.315 (0.957) | 0.474 (0.658) |
| $I/\sigma I$ † | 4.9 (1.5) | 2.8 (1.3) |
| $CC_{1/2}$ † | 0.976 (0.478) | 0.948 (0.346) |
| Completeness (%) | 72.8 (62.4) | 73.3 (42.8) |
| Redundancy | 7.0 (4.6) | 4.9 (1.9) |
| **Refinement** |  |  |
| Resolution (Å) | 29.18-2.50 | 35.89-2.30 |
| No. reflections | 6310 | 8163 |
| $R_{\text{work}} / R_{\text{free}}$ | 0.199/0.253 | 0.234/0.284 |
| No. atoms |  |  |
| Protein | 2049 | 2049 |
| Ligand/ion | 1 | 18 |
| Water | 10 | 34 |
| $B$-factor (Å<sup>2</sup>) | | |
| Protein | 34.81 | 25.18 |
| Ligand/ion | 28.65 | 12.63 |
| Water | 21.93 | 15.51 |
| R.m.s. deviations |  |  |
| Bond lengths (Å) | 0.002 | 0.002 |
| Bond angles (°) | 0.458 | 0.674 |
| Ramachandran |  |  |
| Favoured (%) | 96.86 | 97.25 |
| Allowed (%) | 3.14 | 2.75 |
| Outliers (%) | 0.00 | 0.00 |
| Clashscore | 3.95 | 4.66 |
| Rotamer outliers (%) | 0.00 | 0.00 |

Values in parentheses are for the highest-resolution shell.
\* Merged data from 16 crystals
† Data were truncated at approximately $I/\sigma I \geq 1.0$ and $CC_{1/2} \geq 0.3$ (Karplus & Diederichs, 2012).

**Figure 1.**
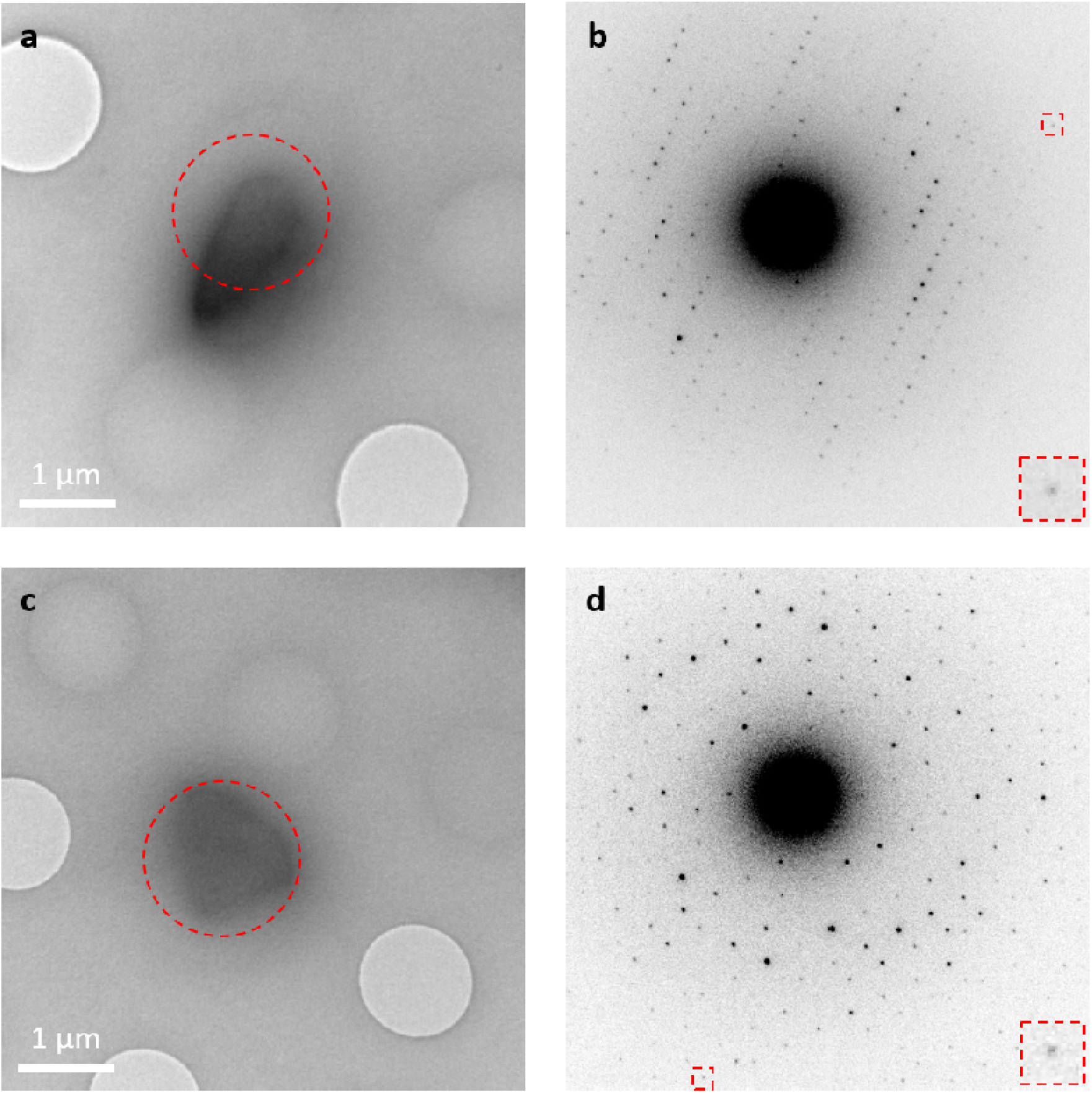
Hydrated 3D microcrystals in a thin layer of vitrified ice. **a.** Micrograph of a native HCA II microcrystal taken at +20.0° tilt. **b.** Electron diffraction pattern of native HCA II recorded over 0.68° tilt at a dose of 0.15 e^−^.Å^−2^. Inset shows a Bragg spot at 2.5 Å resolution. **c.** Micrograph of an inhibitor-bound HCA II:AZM complex microcrystal taken at 0.0° tilt. **d.** Electron diffraction pattern of HCA II:AZM recorded over 0.68° tilt at a dose of 0.15 e^−^.Å^−2^. Inset shows a Bragg spot at 2.3Å resolution. Red circles indicate the area of 1.5 μm in diameter selected for MicroED data collection (selected area electron diffraction mode).

### Structure solution and refinement

The structures of native HCA II and HCA II:AZM complex were phased by molecular replacement, using an apo structure of HCA II with all ligands and solvent removed as search model (Sippel *et al.*, 2009). A well-contrasting single solution was found in space group *P*2_1_ using maximum-likelihood molecular replacement in *Phaser* (McCoy *et al.*, 2007).

Following rigid body refinement of the molecular replacement solution, the resulting electrostatic potential map for the native structure shows clear difference potential indicating the presence of the Zn^2+^ metal cofactor (Fig. 2a). The structure was inspected and modeled using *Coot* (Emsley *et al.*, 2010), fitting the zinc metal cofactor coordinated to the three active site histidine residues (His94, His96, and His119), and refined against the MicroED data (Fig. 2b).

**Figure 2.**
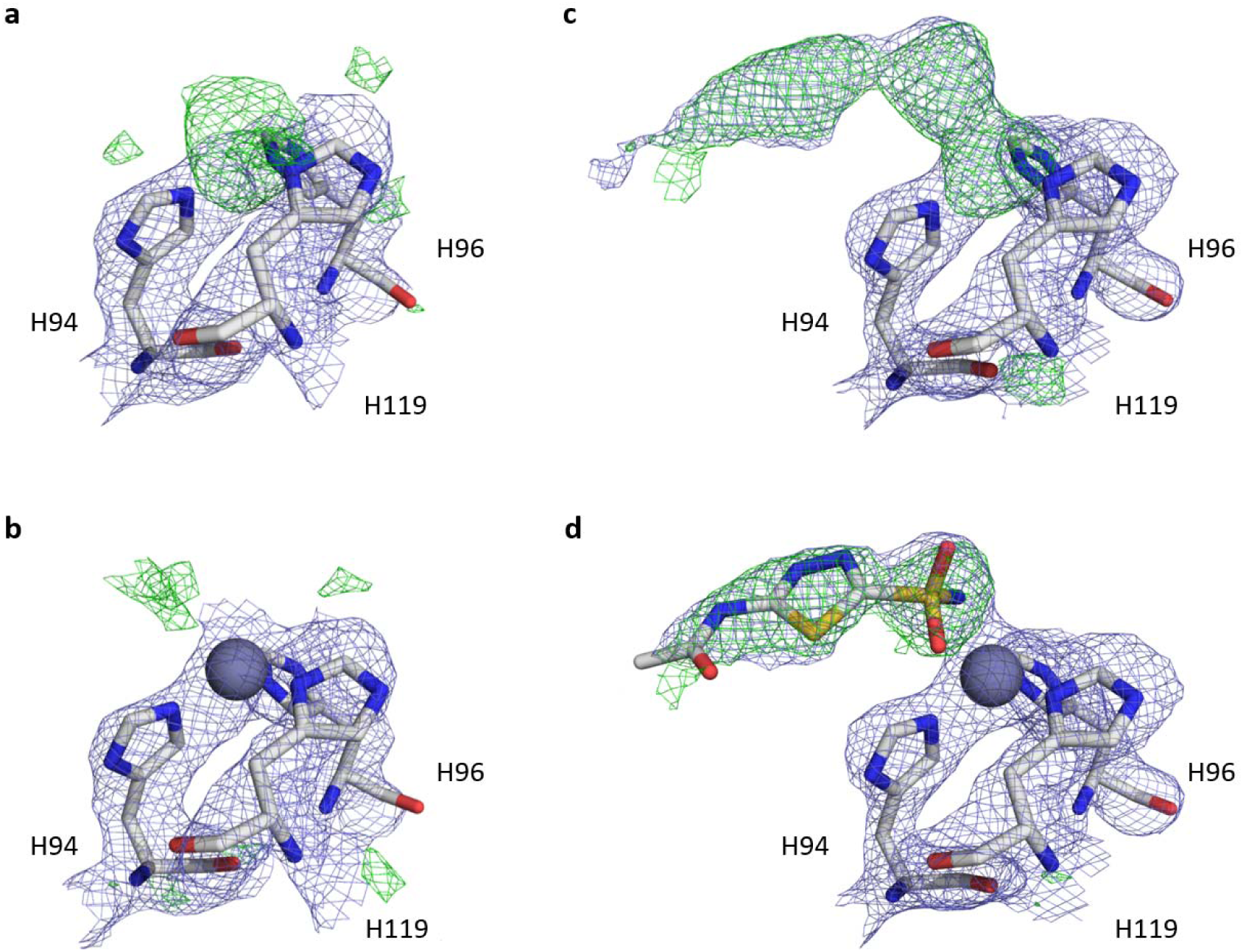
Structure solution and ligand fitting. Electrostatic potential maps displayed for residues of the HCA II active site for the native structure (a,b) and the AZM inhibitor bound model (c,d). **a.** Initial map after rigid body refinement of the molecular replacement solution for the native model, showing clear difference density indicative of the Zn^2+^ metal cofactor. **b.** Refined native model after main chain rebuilding and fitting the metal cofactor in the difference map. **c.** Initial map after rigid body refinement of the molecular replacement solution for the ligand-bound model, showing clear difference density indicative of the Zn^2+^ metal cofactor and AZM inhibitor. **d.** The difference map after rebuilding of the main chain and fitting of the metal was used for placing the drug inhibitor, the found and fitted ligand is superimposed on the structure model, showing its fit to the mFo-DFc map. Electrostatic potential maps 2mFo-DFc are contoured at 1.2σ, colored in blue, and difference maps mFo-DFc contoured at 2.8σ, colored in green and red for positive and negative peaks, respectively. Only observed data were used and no missing reflections were restored for map calculations. Carbon, nitrogen, oxygen and sulphur atoms are colored grey, blue, red and yellow, respectively. Zinc is shown as a dark grey sphere.

For the inhibitor-bound complex, clear difference potential can be observed for the active-site zinc and the AZM inhibitor at its expected position and orientation where it displaces the solvent and zinc-bound water (Fig. 2c). The calculated mFo-DFc difference potential map was then used to fit the AZM inhibitor using *Coot* (Emsley *et al.*, 2010) at a contour level of 2.8σ (Fig. 2d). A final structural model was obtained after real space refinement of the initial inhibitor fit, fixing several geometry outliers and placing solvent molecules (Table 1).

We confirmed that the observed difference potential map used for fitting the inhibitor is not an artefact of model bias by comparing the structure of native HCA II with the structure of HCA II:AZM. The difference potential map of the native structure is only indicative of the metal cofactor and no significant difference signal is observed for the location where the inhibitor is expected to bind (Fig. 2).

### Inhibitor binding interactions

The drug inhibitor AZM has a high affinity for binding HCA II with a K_i_ of 10 nM (Vullo *et al.*, 2003). Furthermore, the active site of HCA II is well accessible for binding of potential inhibitors that block the catalytic activity of the enzyme. The inhibitor binding interactions of AZM are primarily facilitated by its sulfonamide group that replaces the zinc-bound water in the active site. The MicroED structure of inhibitor-bound HCA II does indeed show an active-site zinc that is tetrahedrally coordinated the -NH group of the sulfonamide inhibitor and three histidine residues (Fig. 3a). The histidine to zinc distance is 2.0 Å, which is consistent with previous observations of 2.0 and 2.2 Å from x-ray (PDB ID 3hs4) (Sippel *et al.*, 2009) and neutron diffraction (PDB ID 4g0c) (Fisher *et al.*, 2012), respectively. The lone pair of the sulfonamide N is coordinated directly to the metal co-factor, at a distance of 2.4 Å. This is comparable to available x-ray and neutron models with a distance of 1.9 Å and 2.4 Å, respectively. Additionally, the hydrogen of the sulfonamide N can act as hydrogen donor to Thr199 (nitrogen to oxygen distance of 2.4 Å) that in turn acts as a donor to Glu106 (oxygen to oxygen distance of 2.7 Å). The *B*-factors for the sulfonamide group are within the same range as those of the more stable side chains of the active site residues (Fig. 3b). The AZM thiadiazole ring and acetamido moiety have higher *B*-factors as these are only interacting via weaker hydrophobic and hydrogen bonding, respectively (Fig. 3b).

**Figure 3.**
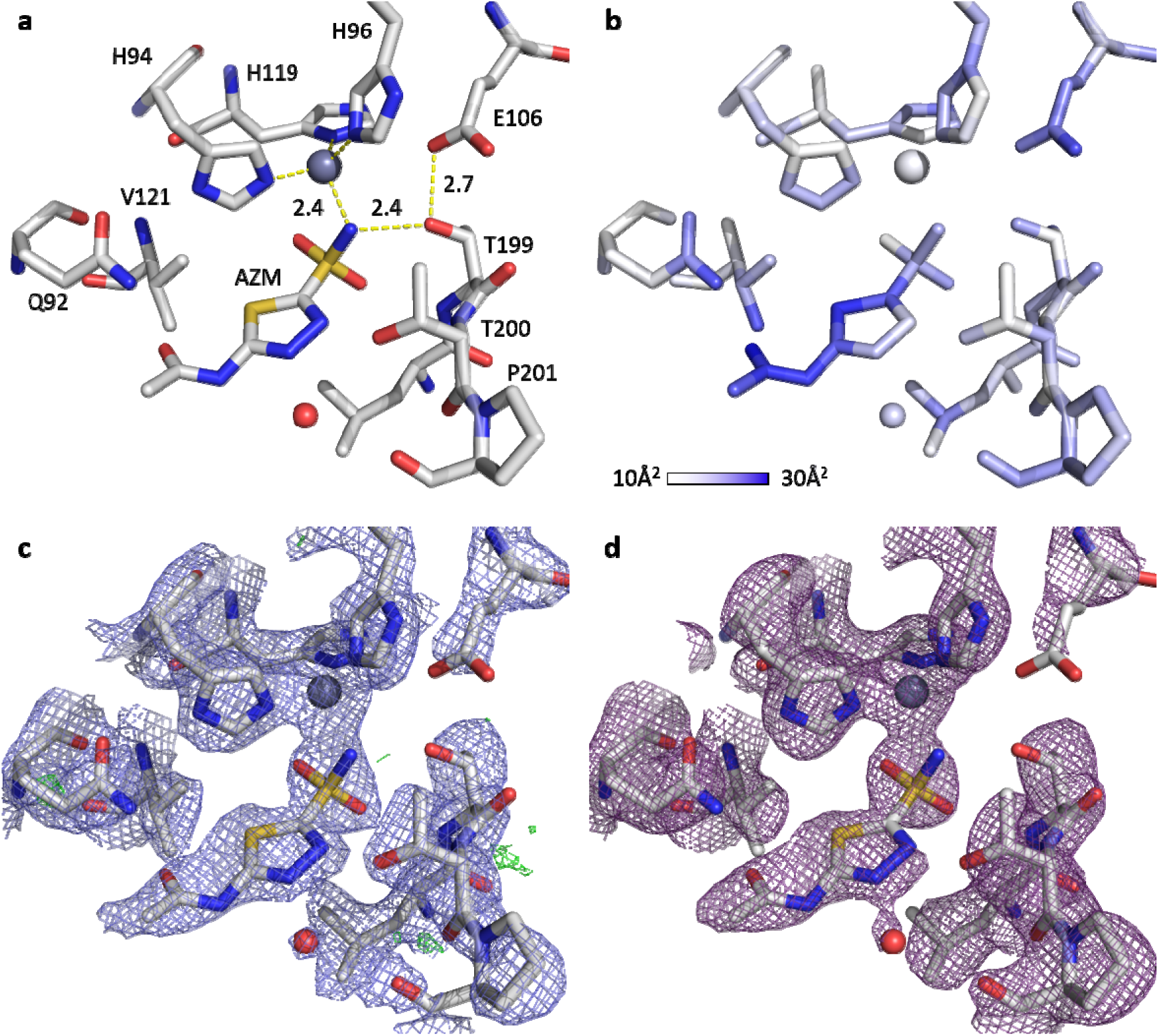
Binding interactions of the AZM inhibitor to the active site of HCA II from the MicroED model. **a.** Interatomic distances measured for binding of the sulfonamide group of AZM, the lone pair of the sulfonamide nitrogen is at a distance of 2.0Å to the active-site zinc, and hydrogen bonding to Thr199 is facilitated at 2.6Å distance measured from nitrogen to oxygen. **b.** *B*-factor values per atom showing a well-ordered sulfonamide group of the inhibitor that is tetrahedrally coordinated to the active site zinc, and interacts via hydrogen bonding with Thr199. **c.** Electrostatic potential map 2mFo-DFc contoured at 1.2σ, colored in blue, and difference potential map mFo-DFc contoured at 2.8σ, colored in green and red for positive and negative peaks, respectively. **d.** Simulated annealing composite omit map is shown for the final model, calculated with sequential 5% fractions of the structure omitted. SA composite omit map is contoured at 1.2σ, colored in magenta. For panels a, c, and d, carbon, nitrogen, oxygen and sulphur atoms are colored grey, blue, red and yellow, respectively. Zinc is shown as a dark grey sphere, water shown as a red sphere. For panels c and d, only observed data were used and no missing reflections were restored for map calculations.

### Structure validation

The refined structural model of the HCA II:AZM shows all features of the active site are well resolved from the electrostatic potential map (Fig. 3c). To validate the structural model with the bound inhibitor, a simulated annealing (SA) composite omit map was calculated, covering the entire unit cell. The SA composite omit map agrees well with the interpretation of the inhibitor-bound protein model (Fig. 3d).

### Comparing MicroED data to x-ray and neutron data

We compare our model and electrostatic potential map against a previously solved structure of the same complex from joint refinement of x-ray and neutron diffraction data (PDB ID 4g0c) (Fisher *et al.*, 2012). The coordination of the metal cofactor and inhibitor in the active site in our model is highly similar to the x-ray and neutron models (Fig. 4). Although the MicroED data have lower resolution (2.3 Å) and completeness (73%) used in refinement compared to those of the x-ray diffraction data (1.6 Å and 94%), our electrostatic potential map can still show all important features, such as well-resolved side chain density and the coordination of the inhibitor bound to the active site (Fig. 4). This is similar to what could be expected from an electron density map using x-ray diffraction data at similar resolution and completeness. The nuclear density map from joint refinement at slightly higher resolution (2.0 Å) and completeness (86%) is less well resolved than our electrostatic potential map, but does provide information about the position of hydrogen atoms in the active site that are difficult to resolve from x-ray data alone.

**Figure 4.**
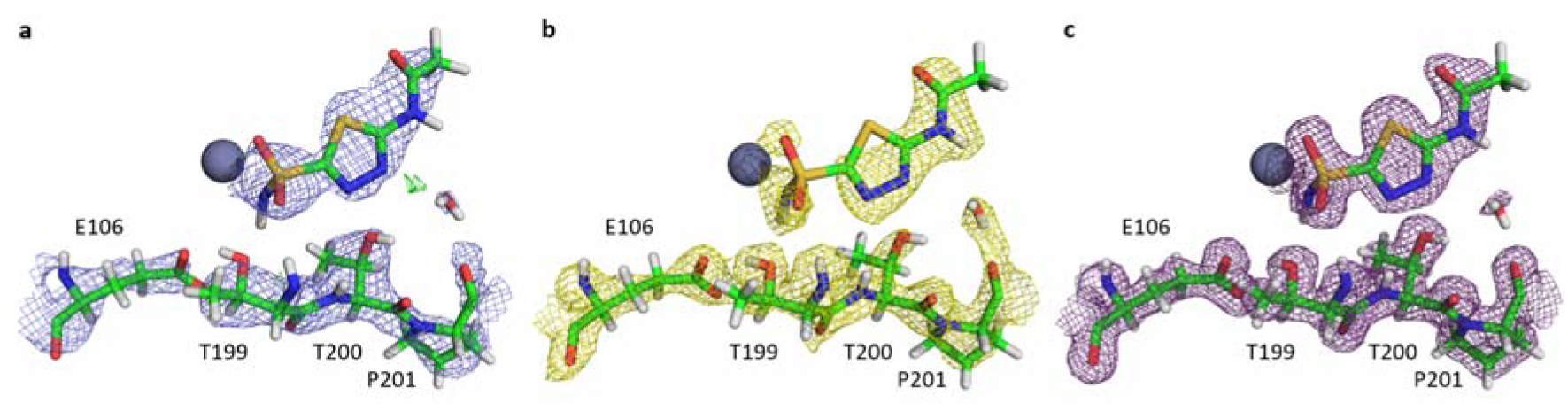
Comparison of the electrostatic potential map from the MicroED model with the electron and nuclear density maps of HCA II with AZM bound (PDB ID 4g0c) (Fisher *et al.*, 2012). **a**. Electrostatic potential map from MicroED data is shown in blue, contoured at 1.5σ (2.3Å resolution, 73.3% completeness). **b.** Nuclear density map is shown in yellow, contoured at 1.5σ (2.0Å resolution, 85.7% completeness). **c.** Electron density map is shown in magenta, contoured at 2.0 σ (1.6Å resolution, 93.6% completeness). Only observed data were used and no missing reflections were restored for map calculations. Hydrogen, carbon, nitrogen, oxygen and sulphur atoms are colored grey, green, blue, red and yellow, respectively. Zinc is shown as a dark grey sphere.

## Discussion

We show that MicroED data can effectively be used for visualizing protein-inhibitor binding interactions by determining the structure of HCA II in complex with clinical drug inhibitor AZM (Fig. 2). The electrostatic potential map was of sufficient quality to allow accurate model building to resolve inhibitor binding. We confirmed the correct interpretation for modelling the structure with the bound inhibitor by analyzing the distances for binding interactions and evaluating our model and data against previously determined structures of the same complex (Fig. 3). Since the structure was phased by molecular replacement, and as data completeness is limited, the structure may be biased by the search model. We present simulated annealing composite omit maps that agree well with the HCA II:AZM model, showing well-defined density for the main chain, side-chains and inhibitor. No missing reflections were filled in for map calculations from weighted estimates of calculated structure factors.

The results presented here indicate MicroED has the potential to resolve inhibitor binding and may play a significant role in future drug discovery and fragment screening. At a resolution of 2.3Å, the data are potentially also suitable for fragment-based screening to identify potential protein inhibitors (Collins *et al.*, 2018). Previous studies could not unambiguously resolve inhibitor-binding interactions based on MicroED data alone for the HIV-1 viral protease and an R2lox metalloenzyme at 2.9 and 3.0Å resolution, respectively (Purdy *et al.*, 2018; Xu *et al.*, 2019). Here we studied the binding of the clinical drug inhibitor AZM that has high affinity to a well accessible active site pocket. A brief soaking of 20 minutes with the dissolved inhibitor proved to be sufficient to observe the inhibitor binding.

MicroED data were efficiently collected from hydrated 3D microcrystals using the rotation method on a conventional transmission electron microscope (TEM), enabling in-house diffraction experiments on a home source for screening and structure determination of biological samples. MicroED thus can complement existing methods in macromolecular crystallography when size requirements dictate the use of larger crystals. In addition, working with protein microcrystals may have several advantages such as less sample material required and fast diffusion of ligand. Another potential benefit of MicroED compared to x-ray diffraction is the possibility to refine atomic charges, exploring the charged state of protein-ligand binding interactions (Yonekura *et al.*, 2015; Yonekura & Maki-Yonekura, 2016). We anticipate that with future hardware and software development optimized towards automated and high-throughput data collection and processing (Cichocka *et al.*, 2018), MicroED can become even more competitive with the ease and speed of synchrotron data collection and fragment screening. Coupled with the potential to refine atomic charges, MicroED may reveal novel details of the underlying molecular mechanisms that mediate inhibitor binding.

## Methods

### Protein expression and purification

HCA II was expressed in BL21 DE3 pLysS *E. coli* cells that were transformed with an expression plasmid based on pET31F1 coding for protein with UniProt accession code P00918(Tanhauser *et al.*, 1992). The cells were grown in shaker incubators at 180 rpm and 37° C in LB media supplemented with 0.1 mg/ml ampicillin. Once the cells reached an optical density of ~1 at 600nm, protein expression was induced with the addition of 1 mM isopropyl β-D-1-thiogalactopyranosid (IPTG) and 1 mM ZnSO_4_. Protein expression was allowed to continue for 3h. At this point the cells were harvested by centrifugation in a JLA 8.1 rotor (Beckman) at ~6800 x *g* for 20 min and frozen at −20 °C. Cell were resuspended in buffer A (0.2 M Na_2_SO_4_, 100 mM Tris pH 9.0) and lysed by addition of lysozyme while stirring in the cold room for 3-4 hours. The cell lysate was centrifuged in a JA-25.50 rotor (Beckman) at ~18000 x *g* for 60 minutes at 4 ºC and the supernatant containing the soluble cellular fraction was loaded onto affinity resin (*para*-aminomethylbenzensulfonamide, Sigma Aldrich A0796). Unbound protein and nucleic acids were removed by repeated wash steps with buffer A, followed by buffer B (0.2 M Na_2_SO_4_, 100 mM Tris pH 7.0). Bound HCA II was eluted with 50 mM Tris pH 7.8 and 0.4 M NaN_3_. The eluted protein was concentrated with Amicon Ultra concentration devices (10 kDa molecular weight cut off), purified and buffer exchanged by size exclusion chromatography on HiLoad 26/600 Superdex 75 (GE Healthcare Life Sciences) in 50 mM Tris pH 8.5. Fractions were collected and analyzed by sodium dodecyl sulfate-polyacrylamide gel electrophoresis (SDS-PAGE) to assess protein purity and homogeneity prior to crystallization. Eluted fractions containing HCA II were pooled and concentrated to ~20 mg/ml by using Amicon Ultra Centrifugal Filter Units (Merck) with a molecular weight cut-off 10 kDa.

### Crystallization

Crystals of HCA II were grown in sitting drop vapour diffusion set-ups using microbridges (Hampton Research) and 24-well Linbro crystallization plates (Hampton Research). The drops were prepared by mixing 10 µl protein solution (20 mg/ml) with 10 µl precipitant solution (2.8 M (NH_4_)_2_SO_4_, 0.1 M Tris pH 8.5) and were equilibrated against 1 ml reservoir solution of the precipitant. Plates were incubated at 20 ºC and crystals appeared in 2-3 days. Crystals were harvested by crushing and repeated pipetting into 1.5 ml Eppendorf tubes to collect slurries of small crystals in mother liquor and stored at 20 ºC until cryo-grid preparation.

### Sample preparation

Complexes of HCA II with inhibitor acetazolamide (Sigma Aldrich, A6011) were prepared by soaking the crystal slurries with inhibitor solubilized in dimethyl sulfoxide (DMSO) for 20 minutes, with a final concentration of 0.5 mM HCA II and 4.5 mM AZM. Grids were prepared by pipetting 3 µl on a QUANTIFOIL 1.2/1.3 (300 mesh) Cu holy carbon TEM grid. Excess liquid was removed via pressure assisted backside-blotting using Preassis (Zhao *et al.*, 2019). The grid was vitrified manually by flash-cooling in liquid ethane. The sample was transferred to a Gatan 914 cryo-transfer holder.

### Data acquisition

Microcrystal electron diffraction data were collected on a JEOL JEM-2100 (LaB6 filament) transmission electron microscope (TEM) operated at 200 kV equipped with a Timepix hybrid pixel detector (Amsterdam Scientific Instruments). Grids were screened for suitable microcrystals in defocused diffraction mode, and diffraction data were collected from an area of 1.5 μm diameter defined by a selected area aperture using the *Instamatic* software interface (Cichocka *et al.*, 2018). The effective sample-to-detector distance was 1481.74 mm. Data were collected using continuous rotation with an angular increment of 0.68° and an exposure time of 1.5 seconds per frame. Individual crystal datasets were typically collected over a tilt range of on average 10-30 degrees, with an acquisition time of 22-66 seconds per crystal dataset. The total range of tilt angles covered during data collection from several crystals was −60 to +60°. The electron dose rate applied during data collection was approximately 0.1 e^−^/Å^2^/s, and the total exposure dose for each crystal dataset was within 2.0-6.0 e^−^/Å^2^.

### Data processing

Data were integrated using *XDS* (Kabsch, 2010). Reflections from different crystal datasets were merged in *XSCALE* using a weighted average of the unit cell parameters obtained from individual crystal processing. The data were analyzed and merged using *AIMLESS* (Evans, 2006). Data were truncated at approximately *I* / σ*I ≥* 1.0 and CC_1/2_ ≥ 0.3 (Karplus & Diederichs, 2012).

**Structure solution.** A search model of the apo structure with the metal cofactor, ligands and waters removed was generated from a high resolution x-ray model (PDB ID 3hs4) (Sippel *et al.*, 2009) using *Sculptor* (Adams *et al.*, 2002). The structure was solved using maximum-likelihood molecular replacement in *Phaser* (McCoy *et al.*, 2007) using the MicroED reflection intensities. For both native and ligand bound data a well-contrasting single solution was found with respectively LLG=1895, TFZ=18.6, and LLG=2247, TFZ=19.7.

### Model building and refinement

Starting electrostatic potential maps were generated from the molecular replacement solution using rigid body refinement in *phenix.refine* (Afonine *et al.*, 2012). The starting model and maps were used for remodeling several sidechains and placing the metal co-factor Zn^2+^ using *Coot* (Emsley *et al.*, 2010). After restrained reciprocal space refinement in *phenix.refine*, the resulting mFo-DFc difference potential map at 2.8σ was used to fit the ligand using *Coot* (Emsley *et al.*, 2010). The fit of the ligand was optimized by real space refinement in *Coot*. A clear blob of difference density was observed in the solvent region, too small to be water but appropriate for DMSO that was used as solvent to solubilize the ligand. The DMSO solvent molecule was fitted using *Coot* (Emsley *et al.*, 2010). Its position is in agreement with the neutron structure (PDB ID 4g0c) (Fisher *et al.*, 2012) where the space is occupied by 3 waters, and with a glycerol (GOL) solvent molecule occupying the same region in the high-resolution x-ray model (PDB ID 3hs4) (Sippel *et al.*, 2009). The MicroED model of HCA II:AZM was then completed using restrained reciprocal space refinement and automatically placing waters with *phenix.refine*. All reciprocal space refinement was preformed using a test set representing 5% of all reflections, atomic scattering factors for electrons, automatic weighting of the experimental data to stereochemistry and atomic displacement parameter terms, and individual *B*-factor refinement (Afonine *et al.*, 2012).

### Validation

The geometry of the native and ligand bound structural models was validated using *MolProbity* (Chen *et al.*, 2010) (Table 1). A simulated annealing (SA) composite omit map was calculated using *phenix.composite_omit_map* (Afonine *et al.*, 2012) by sequentially omitting 5% fractions of the structure. No missing reflections were filled in for map calculations. The MicroED model of the native structure shows a backbone Cα r.m.s deviation of 0.25 Å with a structure from joint refinement against neutron and x-ray diffraction data (PBD ID 3tmj) (Zöe Fisher *et al.*, 2010), and 0.24 Å with an x-ray structure of native HCA II (PBD ID 3ks3) (Avvaru *et al.*, 2010). The inhibitor bound MicroED model shows a backbone Cα r.m.s deviation of 0.24 Å with a previously determined model from joint x-ray and neutron refinement (PDB ID 4g0c) (Fisher *et al.*, 2012), and of 0.15 Å with a high-resolution x-ray model of the same inhibitor bound complex (PDB ID 3hs4) (Sippel *et al.*, 2009), that was used as search model for molecular replacement. Root-mean-square deviation values between structural models were calculated by the secondary-structure matching (SSM) tool (Krissinel & Henrick, 2004).

### Figures

Figures 2-5 were prepared using the PyMOL Molecular Graphics System, version 2.2.3 Schrödinger, LLC.

## Acknowledgements

The project is supported by the Swedish Research Council (2017-05333, H.X.; 2019-00815, X.Z.), the Knut and Alice Wallenberg Foundation (2018.0237, X.Z.), and the Science for Life Laboratory through the technique development grant (MicroED@SciLifeLab)

## Author contributions

M.T.B.C. contributed to sample preparation, electron diffraction data collection, data analysis, structure determination, manuscript writing and figure making. S.Z.F. contributed to project design, crystal growth, sample preparation, and manuscript writing. M.C contributed to data analysis, X.Z. contributed to conception and manuscript writing. H.X. contributed to project design, conception, sample preparation, electron diffraction data collection, data analysis, manuscript writing and figure making.

